# Stem-loops direct precise processing of 3′ UTR-derived small RNA MicL

**DOI:** 10.1101/341578

**Authors:** Taylor B Updegrove, Andrew B Kouse, Katarzyna J Bandyra, Gisela Storz

**Author notes:** These authors contributed equally to this work (Taylor B Updegrove, Andrew B Kouse). Present address: Laboratory of Molecular Biology, National Cancer Institute, Bethesda, MD 20892 USA.

## Abstract

Increasing numbers of 3′UTR-derived small, regulatory RNAs (sRNAs) are being discovered in bacteria, most generated by cleavage from longer transcripts. The enzyme required for these cleavages has been reported to be RNase E, the major endoribonuclease in enterica bacteria. Previous studies investigating RNase E have come to a range of different conclusions regarding the determinants for RNase E processing. To understand the sequence and structure determinants for the precise processing of the 3′ UTR-derived sRNAs, we examined the cleavage of multiple mutant and chimeric derivatives of the 3′ UTR-derived MicL sRNA in vivo and in vitro. Our results revealed that tandem stem-loops 3′ to the cleavage site define optimal, correctly-positioned cleavage of MicL and likely other similar sRNAs. Moreover, our assays of MicL, ArcZ and CpxQ showed that sRNAs exhibit differential sensitivity to RNase E, likely a consequence of a hierarchy of sRNA features recognized by the endonuclease.

## Introduction

While bacterial small RNAs (sRNAs) that act by limited base pairing to increase or repress synthesis from target mRNAs initially were found to be encoded as independent genes in intergenic regions, more and more base pairing sRNAs derived from the 3′UTR sequences of mRNAs are being discovered (Chao *et al*, 2012; Chao & Vogel, 2016; Guo *et al*, 2014; Kim *et al*, 2014; Miyakoshi *et al*, 2015). Some of these sRNAs, such as *Salmonella enterica* CpxQ, arise from the cleavage of the longer mRNA transcript, while others, such as *Escherichia coli* MicL, are transcribed from promoters internal to the protein coding sequence. Even the sRNAs transcribed from the internal promoters can be cleaved to give rise to a shorter product, which, in the example of MicL, contains the region for target base pairing (Guo *et al*, 2014).

The cleavage of the 3′UTR-derived sRNAs characterized thus far has been observed to be well defined, often at or very near the stop codon of the corresponding upstream gene, raising the question of how this specific cleavage occurs. For the *S. enterica* CpxQ sRNA, which is cleaved from the *cpxP* mRNA, the cleavage was found to be dependent on the conserved endoribonuclease RNase E (Chao & Vogel, 2016). In addition, a recent RNA-Seq analysis comparing the 5′ ends of transcripts with and without inactivation of a temperature-sensitive RNase E mutant in *S. enterica* indicates that the majority of 3′UTR-derived sRNAs are generated by RNase E (Chao *et al*, 2017).

Tetrameric RNase E (encoded by *rne*), the major endoribonuclease in enteric bacteria, forms the core of the degradosome (reviewed in (Bandyra *et al*, 2013; Mackie, 2013)). Given the central role of RNase E and the degradosome in the processing of mRNA, rRNA, tRNA, and sRNA (reviewed in (Mohanty & Kushner, 2016)), the determinants of RNase E-dependent cleavage have been studied extensively. Multiple studies have shown that the site of cleavage generally is single stranded and AU-rich (Chao *et al*, 2017; Del Campo *et al*, 2015). Initial characterization of the RNase E cleavage site in the bacteriophage T4 mRNA led to a proposed recognition sequence of (G,A)AUU(U,A) (Ehretsmann *et al*, 1992). In vitro assays examining the cleavage of poly(A) or poly(U) oligonucleotides with substitutions at specific positions further showed that while A- or U-rich sequences are uniformly cleaved, specific nucleotides near the cleavage site impact the position of cleavage (Kaberdin, 2003; McDowall *et al*, 1994). The recent genome-wide analysis of RNase E cleavage sites in *S. enterica* further led to the proposal of a RN|WUU core motif (R = A or G, N = any nucleotide and W = A or U), in which the location of a uridine residue two nucleotides downstream of the cleavage site is most critical (Chao *et al*, 2017).

Considering the degenerate nature of the proposed cleavage motifs, it is not surprising that additional factors have been proposed to impact RNase E-dependent cleavage including the status of the 5′ nucleotide, secondary structure and proteins bound to the RNA. A number of studies have shown that a 5′ monophosphate stimulates RNase E-dependent cleavage sites near this end (Deana *et al*, 2008). However, transcriptome-wide comparisons of total RNA in the presence and absence of the RppH RNA pyrophosphohydrolase, which generates the 5′ monophosphate ends showed that a large percentage of cleavage sites are not impacted by the 5′ end but rather are “direct entry” sites for RNase E (Clarke *et al*, 2014). This study also suggested that the presence of multiple single stranded regions enhanced direct entry by RNase E. Another genome-wide characterization of the secondary structure of the *E. coli* transcriptome via parallel analysis of RNA structure (PARS) coupled to deep sequencing revealed sequences 5′ to 1,800 known RNase E cleavage sites were significantly structured (Del Campo *et al*, 2015). For at least one RNA, the structure was shown to be critical for RNase E cleavage (Ehretsmann *et al*, 1992). Finally, the binding of proteins can impact the position and extent of cleavage both positively and negatively. This is illustrated by RNase E cleavage of the GlmZ sRNA, which is blocked by the binding of the Hfq RNA chaperone protein but is stimulated by the binding of the RapZ protein (Göpel *et al,* 2016). In this example, it was found that RNase E cleaves at a site 6 or 7 nucleotides downstream of the stem-loop RapZ binding site, and conversion of the single stranded region from AU rich to GC rich did not prevent RNase E cleavage, while removal of the upstream stem-loop binding site did.

In contrast to the *S. enterica* CpxQ sRNA, the levels of the truncated *E. coli* MicL sRNA were not strongly reduced in a temperature-sensitive *rne* mutant strain, though levels of the full length transcript increased (Guo *et al*, 2014). Given the prevalence of 3′UTR-derived sRNAs, the precise cleavage observed, and our previous, ambiguous results regarding the ribonucleases acting on the 3′UTR-derived MicL sRNA, we set out to define the determinants for MicL cleavage.

## Results

### ArcZ, CpxQ and MicL are differentially sensitive to the *rne-3071* allele

RNase E is capable of cleaving hundreds of RNAs and has been shown to play the most prominent role in sRNA cleavage in *E. coli* (reviewed in (Mohanty & Kushner, 2016)). The enzyme is essential though overexpression of RNase G, which can cleave many of the same targets, can rescue an *rne*- strain (Lee *et al*, 2002). To determine the impact of RNase E and RNase G on cleavage of MicL, total RNA was isolated from strains carrying combinations of wild-type and temperature-sensitive *rne-3071* (Ono & Kuwano, 1979) and Δ*rng* alleles. Strains initially grown at 30°C to OD600~1.0 were split and cultured for an additional h at either 30°C or 43.5°C, after which total RNA was extracted.

We first assayed the levels of ArcZ and CpxQ, previously reported to be cleaved by RNase E (Chao *et al*, 2017; Chao & Vogel, 2016). As expected, while the levels of the cleavage products were similar for all of the strains grown at 30°C (Fig 1A, lanes 1-4), the levels of the processed transcripts were significantly reduced in the *rne-3071* single mutant while the levels of the longer, uncleaved transcripts increased at the non-permissive temperature of 43.5°C (Fig 1A, lane 6). The products detected for the Δ*rng* single mutant were similar to the products seen for the wild type strain at 43.5°C (Fig 1A, lanes 5 and 7). In contrast, the cleavage products were almost entirely absent in the *rne-3071* Δ*rng* double mutant (Fig 1A, lane 8).

**Figure 1.**
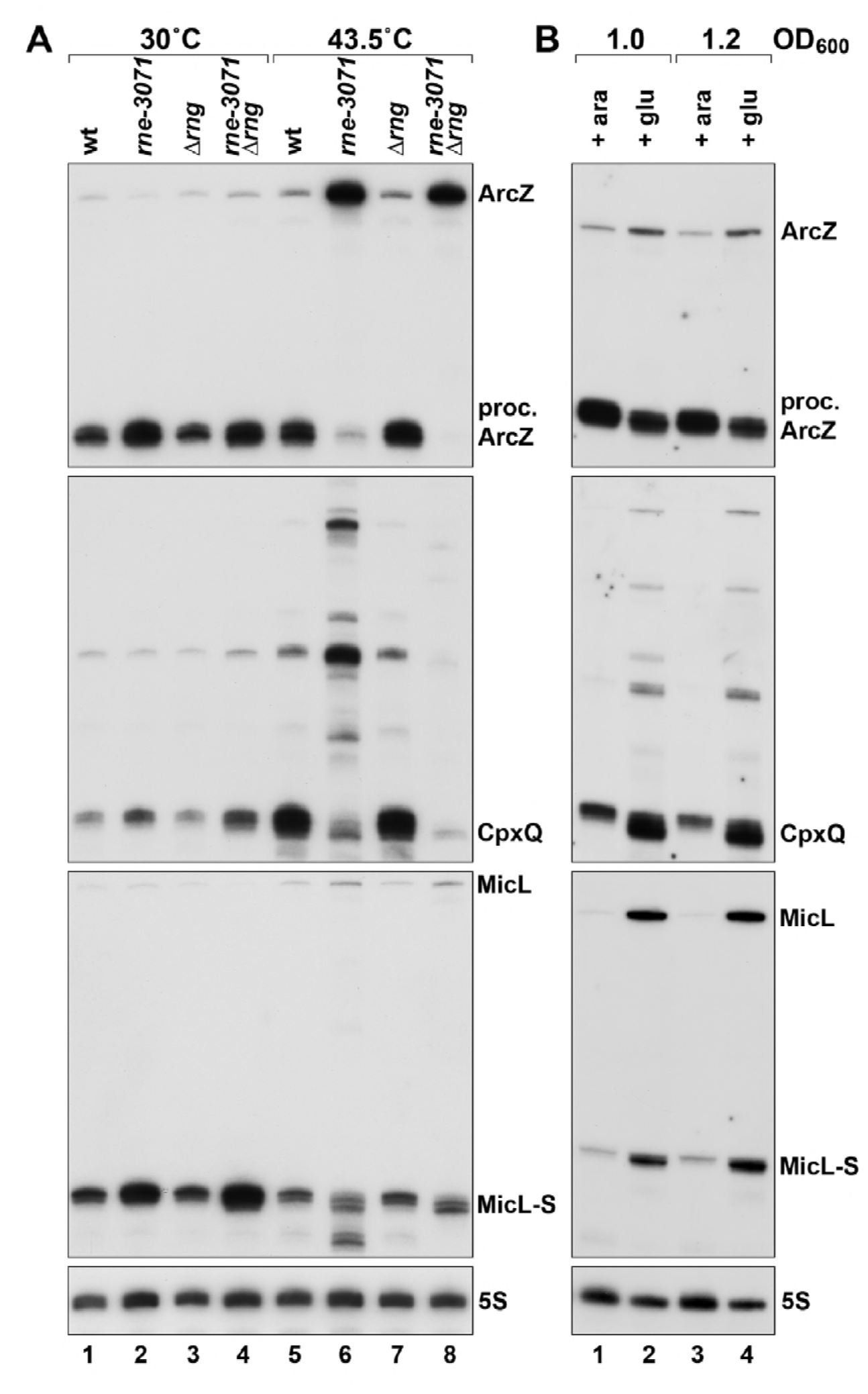
Varied effects of rne-3071, Δ*rng* and RNase E depletion on ArcZ, CpxQ and MicL cleavage. A. Four *E. coli* strains, wild type (EM1279), *rne-*3071 (EM1277), Δ*rng* (EM1279 Δ*rng*) and *rne-*3071 Δ*rng* double mutant (EM1277 Δ*rng*) were cultured in LB at 30°C to an _OD600_ of 1.0. Cultures were then split in half and incubated at either 30°C or 43.5°C for 1 h.
B. *E. coli* KSL2000, with the pBAD-RNE plasmid was cultured in LB arabinose at 37°C to an OD_600_ of 0.5. Cultures were split and half washed and grown in LB arabinose and the other half washed and grown in LB glucose to OD_600_ of 1.0 or 1.2. For both panels, total RNA extracted from the cultures was subjected to northern analysis using probes against the ArcZ, CpxQ, MicL and 5S RNAs.

The effects of the *rne-3071* and *rne-3071* Δ*rng* mutations were somewhat different for MicL when the same northern membrane was probed for this sRNA (Fig 1A). Similarly, low levels of MicL and high levels of cleaved MicL (MicL-S) transcripts were found across each of the 30°C samples (Fig 1A, lanes 1-4). For the cells shifted to 43.5°C, the overall levels of MicL were higher and MicL-S levels were lower. Again, the patterns for wild type and Δ*rng* strains were similar (Fig 1A, lanes 5 and 7). However, unlike for ArcZ and CpxQ, the levels of cleaved MicL-S were not greatly reduced in the *rne-3071* single and *rne-3071* Δ*rng* double mutant strains. The levels of MicL increased, and the pattern of MicL-S was changed somewhat with the detection of one slightly shorter transcript in both mutants as well as somewhat shorter RNase G-dependent products in the *rne-3071* mutant (Fig 1A, lanes 6 and 8).

### ArcZ, CpxQ and MicL are differentially sensitive to RNase E depletion

We also examined the consequences of depleting wild type RNase E in an *rne*- strain with RNase E expressed from a plasmid behind a P_BAD_ promoter (Lee *et al*, 2002). A culture of these cells was grown to mid-exponential phase in LB with arabinose, split and the two halves were washed with either LB with arabinose, which induces expression from the _PBAD_ promoter, or LB with glucose, which represses expression. Total RNA isolated from the two cultures further grown to either OD_600_ ~1.0 or ~1.2 was again probed for the ArcZ, CpxQ and MicL sRNAs (Fig 1B). Full-length and processed ArcZ as well as CpxQ were present at approximately wild type levels for the cells grown with arabinose to induce RNase E (Fig 1B, lanes 1 and 3). In contrast, the levels of MicL-S were reduced suggesting that this sRNA may be more sensitive to elevated RNase E likely expressed from the plasmid. As expected, depletion of RNase E by growth in glucose led to increased levels of full-length and decreased levels of the processed ArcZ though cleavage was not completely abolished (Fig 1B, lanes 2 and 4). Interestingly, the levels of CpxQ, as well as intermediate cleavage products observed with the *rne-3071* strain, all increased for the glucose-grown cultures (Fig 1B, lanes 2 and 4). MicL and MicL-S levels also were both increased with glucose with a greater increase for full-length MicL (Fig 1B, lanes 2 and 4). Thus, as observed for the *rne-3071* strain, the three sRNAs showed differential sensitivity to altered RNase E levels.

### MicL is cleaved by RNase E in vitro

To further test whether RNase E is capable of cleaving MicL at the position observed in vivo, we carried out in vitro cleavage assays with the purified catalytic N-terminal domain (NTD; residues 1-529) of RNase E. As shown in Fig 2, when we incubated full length MicL with RNase E (1- 529) and the RNA chaperone Hfq, we observed faint cleavage of MicL at the position where cleavage is seen in vivo. The levels of this product increased when the RNA was incubated with a mutant RNase E (1-529) (D26N, D28N and D338N) that is more catalytically active than wild-type RNase E (1-529) (Bandyra *et al*, 2018).

**Figure 2.**
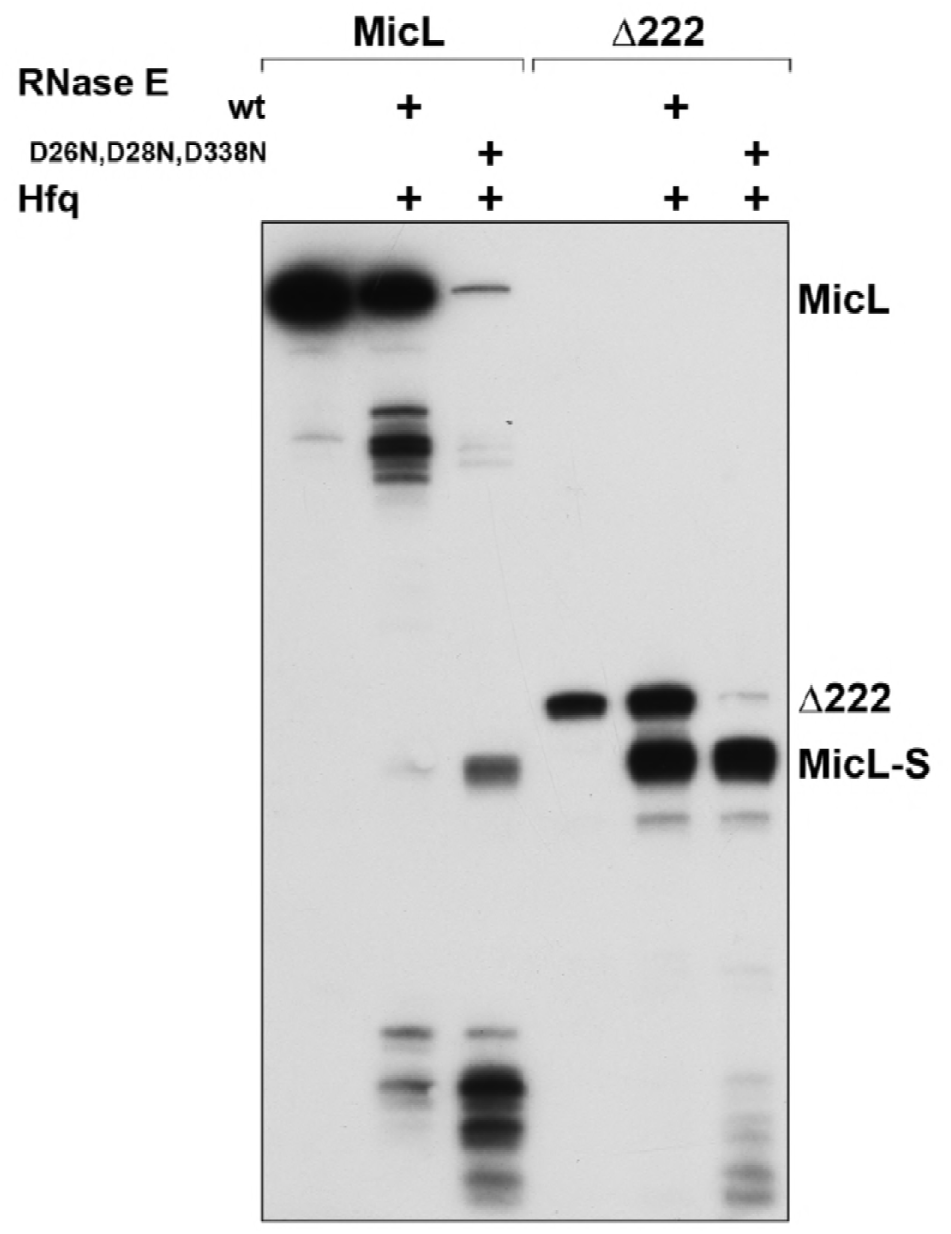
Purified RNase E (1-529) cleaves MicL in vitro. In vitro transcribed full-length MicL (308 nt) and Δ222 (86 nt) with 5′PPP was incubated with purified Hfq and purified wild type RNase E (1-529) or D26N,D28N,D338N mutant RNase E (1- 529) at 30°C for 30 min. The RNA was then subject to northern analysis using an oligonucleotide probe complementary to the 3′-end of MicL.

A truncated version of MicL lacking the first 222 nt, which is processed as efficiently as wild type MicL in vivo (see below), is cleaved robustly by both wild type and mutant RNase E (1-529) (Fig 2). Together the results of the in vivo and in vitro assays indicate that MicL is cleaved by RNase E, perhaps as a highly sensitive substrate that is still processed by low levels of the endonuclease.

### Cleavage of MicL is not affected by sequences at the 5′ end

We next wanted to examine what sequences directed the very specific cleavage of MicL to give MicL-S. To determine if sequences 5′ of the MicL cleavage site are important, we examined the consequences of sequential deletions of this region (Fig 3A and Fig EV1A). The series of 5′ truncations were cloned into the pBRplac* expression plasmid and introduced into a Δ*cutC* strain, which lacks the native *micL* promoter and the region of *micL* overlapping the *cutC* coding sequence (Guo *et al*, 2014). Total RNA isolated from these strains was examined by northern analysis using an oligonucleotide probe complementary to the 3′ end of MicL (Fig 3B and Fig EV1B).

**Figure 3.**
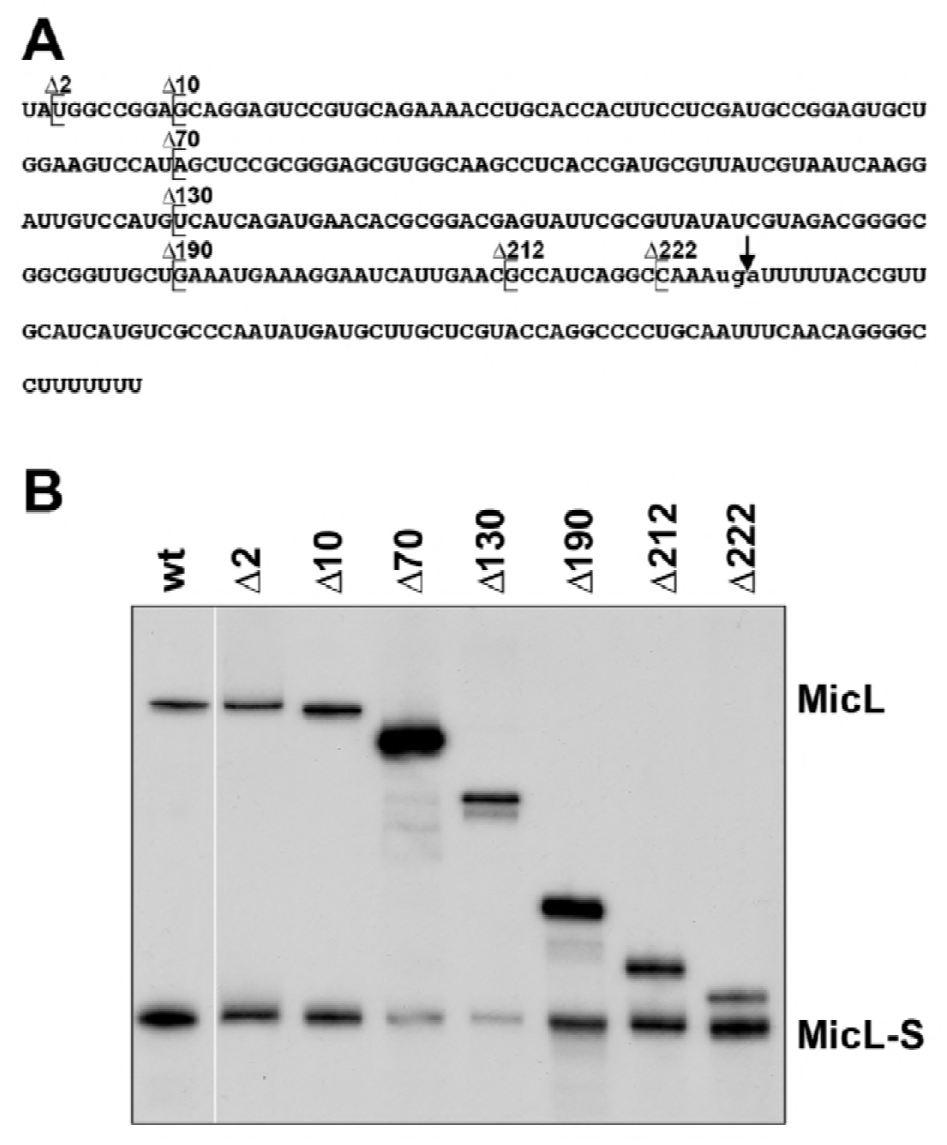
Sequences at the 5′end of cleavage site are not required for MicL cleavage. A. A MicL RNA sequence. Lower case sequence and arrow indicate *cutC* stop codon and RNase E cleavage site, respectively, and brackets denotes the portion of MicL 5’end deleted (as indicated above the sequence) in constructs assayed in Fig 3B.
B. Wild-type (wt) and the indicated MicL mutants with 5’ end deletions were cloned into the pBRplac* plasmid. Total RNA isolated from a Δ*cutC* strain carrying these constructs was subject to northern analysis using an oligonucleotide probe complementary to the 3’ end of MicL.

All of the constructs still gave rise to the MicL-S cleavage product, though the levels were reduced for Δ70, Δ72, Δ82, Δ130, Δ132 and particularly Δ162. Structure predictions suggested that for the constructs with reduced MicL-S levels, the cleavage site might be occluded by the formation of an alternative stem-loop (Fig EV1A). To test this possibility, we mutated residues 177-180 and 204-208 predicted to be involved in this pairing and observed that cleavage is increased (Fig EV1C). Overall, given that the same MicL-S cleavage product was observed for all 19 constructs assayed, we concluded that the sequences 5′ of the cleavage site are not important for specific cleavage, but that, as has also been previously reported for other RNase E substrates (Chao *et al*, 2017; Ehretsmann *et al*, 1992; McDowall *et al*, 1994), the cleavage site must be single stranded. The Δ222 deletion construct, which contained only six nucleotides upstream of the cleavage site but still exhibited robust processing comparable to wild type MicL in vivo (Fig 3B) and in vitro (Fig 2), was chosen as the starting construct for further experiments.

### Sequences surrounding the cleavage site can be promiscuous

Studies in multiple organisms have indicated that RNase E preferentially cleaves at sequences rich in adenine and uridine (Ehretsmann *et al*, 1992). Additionally, a genome-wide analysis led to a proposed RNase E consensus sequence of RN|WUU, with the U at the +2 position being the most critical (Chao *et al*, 2017). The MicL cleavage site of AAAUG|AUUU is a good match to the consensus sequence. To examine the contribution of this sequence to cleavage, three mutants were constructed in which three adjacent residues, AAA preceding the site, UGA overlapping the site and three U residues after the site, were mutated to CCC (Fig 4A), given that C residues were previously shown to inhibit RNase E cleavage, particularly two nucleotides downstream of the cleavage site (Chao *et al*, 2017; Ehretsmann *et al*, 1992). Total RNA isolated from strains expressing these derivatives of Δ222 was subject to northern analysis. As expected, given that the A residues 5′of the cut site did not match the consensus, mutation of these residues did not change cleavage efficiency. Surprisingly however, mutations to the cut site and the U residues 3’ of the cut site also did not greatly affect cleavage efficiency (Fig 4B).

**Figure 4.**
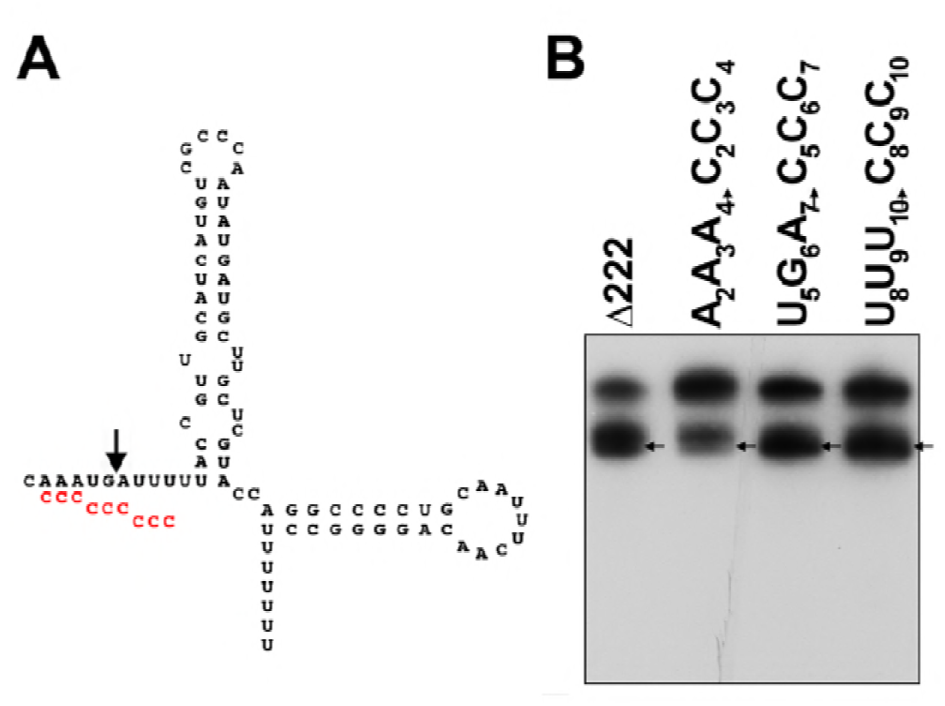
Sequences surrounding the MicL cleavage site can be varied. A. Predicted secondary structure of Δ222 constructs with mutations made in separate constructs indicated in red below the nucleotides which they replace. The arrow indicates site of RNase E cleavage.
B. B The Δ222, A_2_A_3_A_4_ ➜ C_2_C_3_C_4_, U_5_G_6_A_7_➜C_5_C_6_C_7_ and U_8_U_9_U_10_ ➜ C_8_C_9_C_10_ constructs were expressed from the pBRplac* vector in Δ*cutC* strains. Total RNA was extracted from each culture and probed for MicL as in Fig 3B. Arrows denote cleavage products corresponding to the site indicated in Fig 3A.

Similar results were seen for full-length MicL, where a plasmid overexpressing MicL carried mutations disrupting the cleavage site sequence or the flanking single-stranded sequences (Fig EV2). In this case, the UG|A cut site sequence was mutated to GG|A, UC|G, UC|A or GC|C and the flanking 5′ AAA and 3′ UUU sequences (relative to the UG|A cut site) were mutated to UUU and AAA respectively. None of the aforementioned mutations affected cleavage. These data indicate MicL cleavage is not greatly influenced by the sequence directly adjacent to or overlapping the cleavage site.

### Stable 3′ stem-loops are required for efficient cleavage

Having ruled out the sequence 5′ to and overlapping the cleavage site as necessary determinants of MicL cleavage, we next examine the region 3′ of the cleavage site. This region contains two predicted secondary structural elements: stem-loop A (ΔG = −11.7 kcal/mol) and the terminator, stem-loop B (ΔG = −17.2 kcal/mol). Given the presence of stem-loops adjacent to other RNase E cleavage sites in *E. coli* (Del Campo *et al*, 2015), we wondered whether the stem-loops found in the Δ222 derivative might affect its cleavage efficiency. We first tested whether the sequences of the two stem-loops were important by replacing them with stem-loops from Spot 42 sRNA, which have a similar size and stability despite significantly different sequences (Fig 5A). Analysis of total RNA isolated from cells expressing these constructs revealed that the heterologous stem-loops did not affect the location or efficiency of cleavage; the pattern was very similar for Δ222, stemA=Spot 42 (ΔG = −10.8 kcal/mol) and stemB=Spot 42 (ΔG = −21.2 kcal/mol) (Fig 5B) indicating the sequences of the 3′ stem-loops do not impact cleavage.

**Figure 5.**
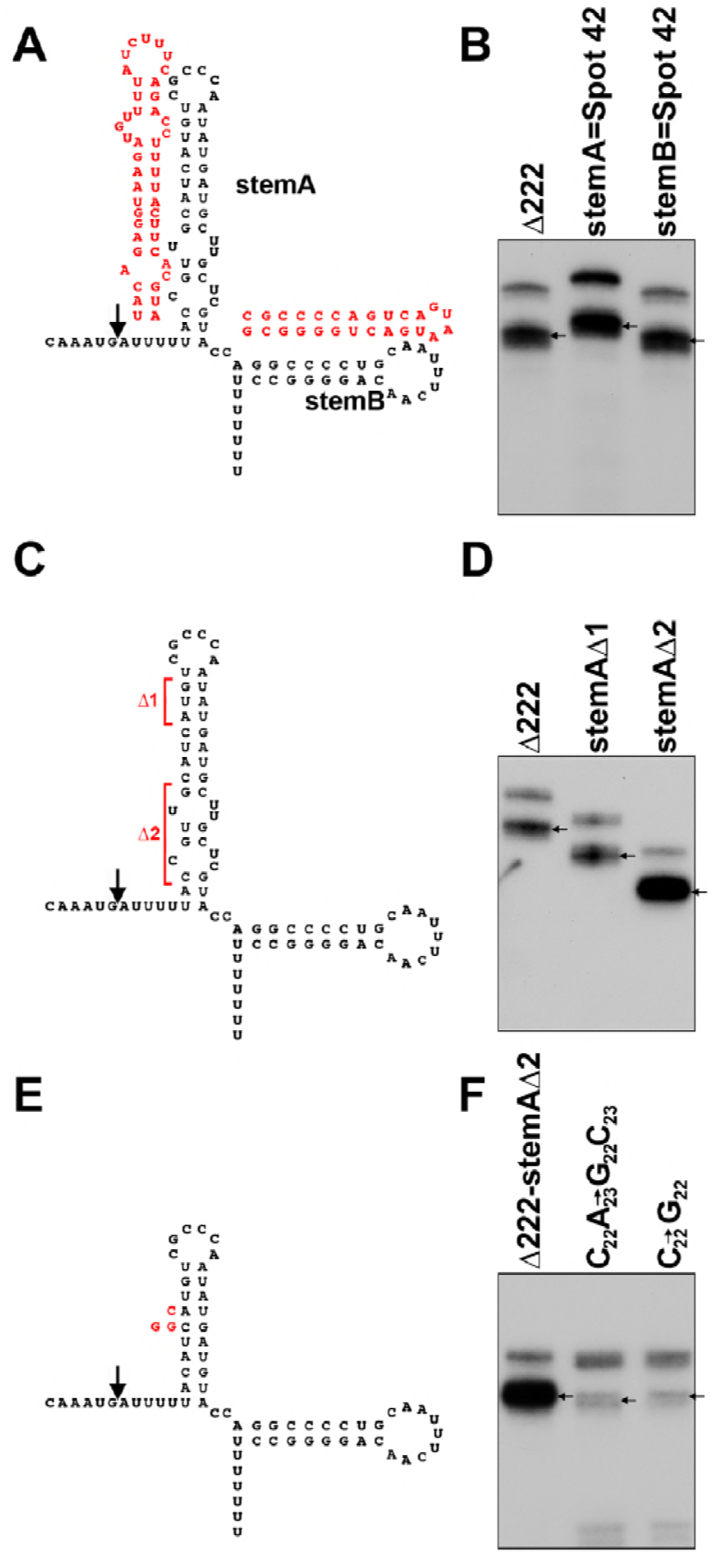
Two stable stem-loops are required for efficient MicL cleavage. A. Predicted secondary structure of Δ222 constructs with stem-loop A and stem-loop B replaced by similar stem-loops from Spot 42. The arrow indicates site of RNase E cleavage.
B. The constructs depicted in Fig 5A were expressed from pBRplac* in Δ*cutC* background. Arrows denote cleavage products corresponding to the site indicated in Fig 5A.
C. Predicted secondary structure of Δ222 constructs with regions of stem A deleted indicated in red brackets. The arrow indicates site of RNase E cleavage.
D. Constructs stemAΔ1 and stemAΔ2 depicted in 5C were expressed from the pBRplac* vector in Δ*cutC* background. Arrows denote cleavage products corresponding to the site indicated in Fig 5C.
E. Predicted secondary structure of Δ222-stemAΔ2 construct with red nucleotides indicating residues changed in the C_22_A_23_ ➜ G_22_C_23_ and C_22_ ➜ G_22_ constructs. The arrow indicates cleavage site.
F. The constructs depicted in Fig 5E were expressed from pBRplac* in Δ*cutC* background. Arrows denote cleavage products corresponding to the site indicated in Fig 5E. For samples in panels B, D, and F, total RNA was extracted and probed for MicL as in Fig 3B.

To further examine whether stem-loop A was important, we shortened this stem in two mutants. In stemAΔ1 (ΔG = −8.7 kcal/mol), the stem was shortened by the removal of three base pairs towards the top of the stem-loop and in stemAΔ2 (ΔG = −13.0 kcal/mol), the bulges in the lower portion of the stem were removed thereby increasing stability (Fig 5C). Again, synthesis from these constructs was examined by northern analysis. Interestingly, while the stemAΔ1 construct showed cleavage similar to the control Δ222 construct, the stemAΔ2 derivative, which is predicted to be more stable than the wild type stemA, gave higher levels of MicL-S (Fig 5D). These data suggest that stem-loop A influences cleavage efficiency, possibly along with contributing to transcript stability.

To further explore the effect of stem A stability on processing, we additionally introduced mutations to destabilize the stem in the context of Δ222 with stemAΔ2; C_22_A_23_➜G_22_C_23_ (ΔG = −4.1 kcal/mol) and C_22_➜G22 (ΔG = −5.8 kcal/mol) (Fig 5E). The transcripts made from these constructs were then compared to the Δ222-stemAΔ2 (ΔG = −13.0 kcal/mol) RNA. Both destabilized constructs showed a decrease in the cleavage product when compared to Δ222-stemAΔ2, further suggesting that the stability of stem-loop A affects cleavage efficiency (Fig 5F).

### Stem-loop A determines position of cleavage

To investigate whether stem-loop A also governs the position of cleavage, we inserted heterologous sequences of ACACAC, UCUCUC or UGUGUG in the single-stranded region between the cleavage site and stem-loop A and examined the influence of the insertions on the cleavage of the Δ222 construct (Fig 6A). Northern analysis of all three mutants expressed from a plasmid (Fig 6B) clearly shows cleavage to generate MicL-S, again suggesting cleavage can occur at a very precise distance of 5 nucleotides from stem-loop A regardless of sequence. We also detect an extra band, particularly for the UCUCUC and UGUGUG mutants that is slightly larger than MicL-S. Interestingly, alternative structures that extend stem-loop A can be predicted for these two constructs.

**Figure 6.**
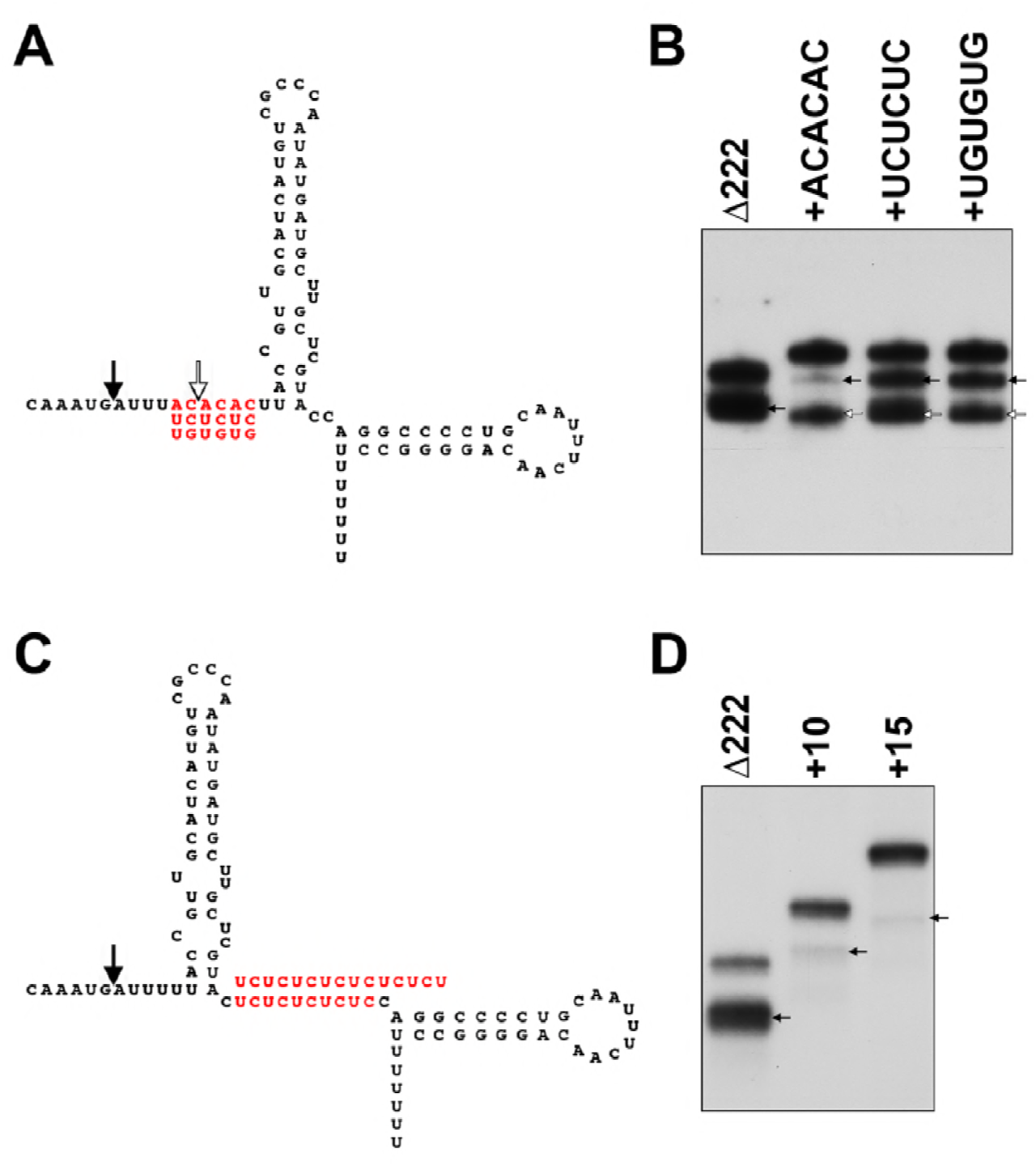
Distances from 5′ stem-loop and between two stem-loops of MicL affect cleavage. A. Predicted secondary structure of Δ222 constructs with insertions made in separate constructs given in red. The black arrow indicates cleavage site in Δ222, and the white arrow indicates the new cleavage site observed.
B. Constructs depicted in Fig 6A were expressed from pBRplac* in Δ*cutC* background. Arrows denote cleavage products corresponding to the sites indicated in Fig 6A.
C. Predicted secondary structure of Δ222 constructs with spacer sequences introduced between stem-loop A and stem-loop B given in red. The arrow indicates cleavage site.
D. Constructs depicted in Fig 6C were expressed from pBRplac* in Δ*cutC* background. Arrows denote cleavage products corresponding to the site indicated in Fig 6C. For samples in panels B and D, total RNA was extracted and probed for MicL as in Fig 3B.

### Distances between two 3′ stem-loops of MicL affect cleavage

We noted that the two stem-loops are directly adjacent to each other and next examined the consequences of altering the distance between the two structures with repeating U and C ‘spacers’ of 10 or 15 nucleotides between stem-loop A and B in the Δ222 context (Fig 6C). Secondary structure predictions indicate the spacer constructs create a single stranded region between the stem-loops and do not alter the stem-loop structures and their stabilities. Again, total RNA isolated from strains carrying the resulting constructs was subjected to northern analysis (Fig 6D). Compared to the Δ222 control, both spacer constructs showed decreased levels of cleavage with less product for the +15 construct than for the +10 construct. These data indicate that two stable stem-loops in close proximity to the RNase E cleavage site are required for efficient MicL cleavage, superseding recognition of a specific sequence.

### Strongest cleavage is observed with two adjacent 3′ stem-loops

To further test whether two stem-loops are needed for efficient cleavage and to examine the influence of these two stem-loops with respect to the distance from and position 5′ or 3′ of the cleavage site, we synthesized a number of synthetic constructs (Fig 7A) starting with the Δ222-stemAΔ2 MicL derivative. For all of the synthetic constructs, the same linear sequence of four repeats of the MicL cleavage site (UGAUU) separated by a CC or UC spacer was flanked by different stem-loops. The repeat sequence was predicted to be single-stranded and not affect the structures of added or altered stem-loops in each of the constructs. Total RNA extracted from the strains harboring the resulting vectors was subject to northern analysis (Fig 7B). For the construct with the repeats followed by stemAΔ2 and the wild type stem-loop B terminator (repeats+stemAΔ2+term, Fig 7B, lane 2), the 95 nt transcript gave the same predominant MicL-S product as the Δ222-stemAΔ2 construct (Fig 7B, lane 1), though one additional minor cleavage product was observed. These data suggest that although additional putative cleavage sites are present in the 5′-single stranded region, cleavage was directed towards the site nearest the two 3′ stem-loops.

**Figure 7.**
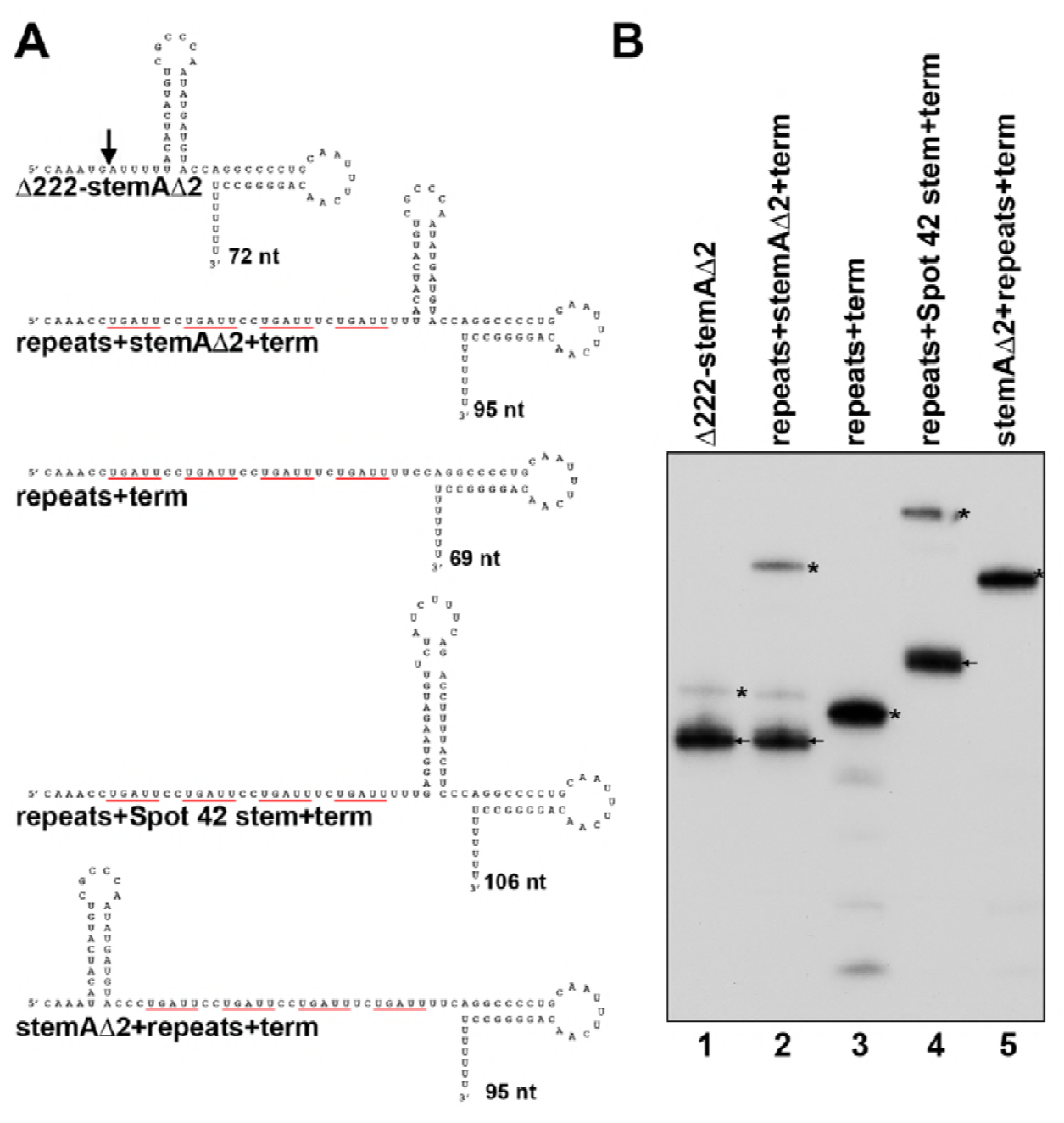
Strongest cleavage is observed with two 3’ stem-loops. A. Predicted secondary structures of constructs with repeat sequences between different stem-loop structures. Secondary structures were predicted using the Mfold software package. The arrow indicates the site of RNase E cleavage. UGAUU repeats are underlined in red.
B. Cleavage is directed to the site nearest to two stable stem-loops. The constructs from Fig 7A were expressed from pBRplac* in Δ*cutC* background. Total RNA was extracted and probed for MicL. Full length products are denoted by asterisks, and prominent cleavage products are denoted by arrows.

The requirement for having two 3′ stem-loops was again tested by deleting stemA (repeats+term). Analysis of the total RNA from the strain carrying this construct showed there was very little cleavage of the 69 nt transcript with just stem-loop B (Fig 7B, lane 3). We suggest that the four minor products observed correspond to cleavage at the four consensus sequences in the linear repeat. Addition of a Spot 42 stem-loop in place of stem A (repeats+Spot 42 stem+term) restored strong cleavage at the position closest to the stem-loops (Fig 7B, lane 4), further indicating that efficient cleavage requires two stable stem-loops, independent of the sequences of the stem-loops.

To test whether stem-loops at the 5′ end of the repeat sequence could also influence cleavage, we examined one additional construct carrying stemAΔ2 upstream and the stem-loop B terminator downstream of the repeat sequence (stemAΔ2+repeats+term). Interestingly, only very faint cleavage products were detected for this construct (Fig 7B, lane 5). Thus, two adjacent stem-loops are required for robust cleavage.

### Mutations in MicL and RNase E decrease MicL cleavage in vitro

Finally, we tested whether the effects of the MicL mutations and insertions similarly impacted cleavage by purified RNase E. In vitro transcribed Δ222, stemAΔ2, +10 and +15 were all incubated with purified RNase E (1-529) and examined by northern analysis. The results of the in vitro experiments are consistent with the in vivo findings with stemAΔ2 showing somewhat increased cleavage and +10 and +15 showing decreased cleavage (Fig 8A).

**Figure 8.**
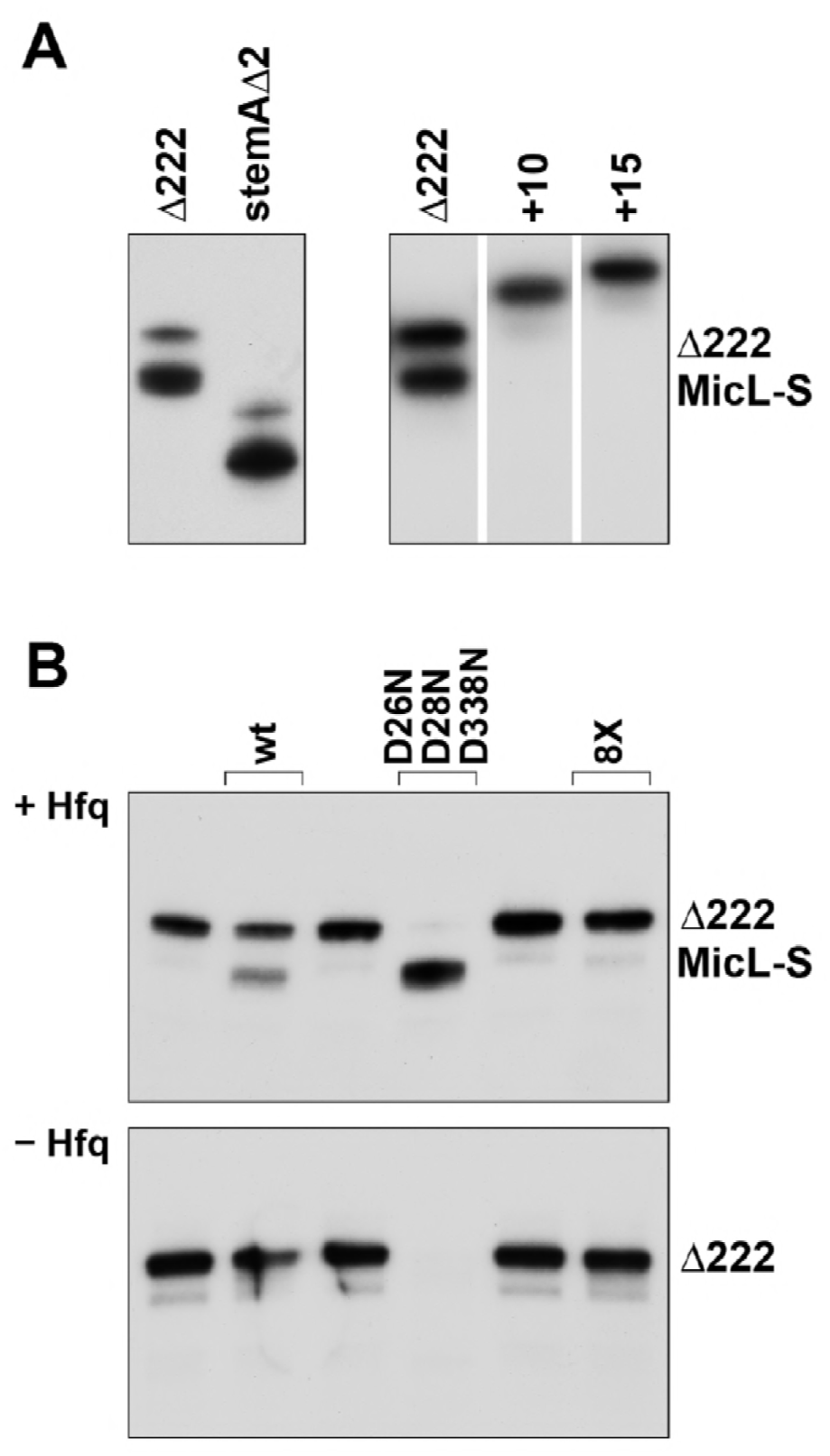
Effects of mutations on MicL cleavage in vitro. A. In vitro transcribed Δ222, Δ222-stemAΔ2, Δ222+10 and Δ222+15 with 5′PPP were incubated with purified Hfq and purified wild type RNase E (1-529) at 30°C for 30 min.
B. In vitro transcribed Δ222 was incubated with and without purified Hfq and purified wild type RNase E (1-529), D26N,D28N,D338N mutant RNase E (1-529) or 8X mutant RNase E (1- 529) at 30°C for 30 min. For both A and B, the RNA was subject to northern analysis using an oligonucleotide probe complementary to the 3′-end of MicL.

Two recent crystal structures of a catalytically inactive RNase E (1-529 D303R D346R) with sRNA fragments (Bandyra *et al*, 2018) have shown that helices of both RprA and SdsR are recognized by RNase E through an interaction with the RNase H like domain and small domain located distal to the catalytic active site of RNase E. The structures led to the identification of eight amino acid residues (R3, Q22, H268, Q270 and Y269 in the RNase H domain and K433, R488 and R490 in the small domain) that when mutated (RNase E 1-529 8X mutant) impaired RNase E binding to and cleavage of RNA substrates with helical elements adjacent to the cleavage site. Among the substrates tested was the 9S rRNA, previously reported to contain structural elements important for RNase E recognition (Cormack & Mackie, 1992). We similarly did not observe cleavage of the 9S rRNA unless the in vitro transcript was treated with calf intestinal phosphatase (CIP) to convert the 5′ triphosphate to monophosphate to allow 5′ end recognition (Fig EV3). Consistent with RNase E recognition of stem-loop structures in MicL, the Δ222 derivative was not cleaved by the 8X RNase E mutant (Fig 8B).

Up to this point, we always included the Hfq protein with RNase E in all our in vitro cleavage assays with MicL. To examine the contribution of the RNA chaperone to the cleavage, we also carried out a set of reactions in which we omitted Hfq (Fig 8B). Interestingly, we no longer detected Δ222 cleavage with wild type RNase E (1-529) while the RNA was completely digested by the hyperactive D26N,D28N,D338N mutant. These results indicate that Hfq may have two roles in promoting MicL cleavage; positioning RNase E to help direct cleavage at a specific position as well as protecting the cleaved product from excess cleavage.

## Discussion

The 308 nt σ^E^-dependent MicL sRNA, transcribed from within the *cutC* gene, was previously reported to be cleaved to give a 80 nt derivative capable of binding Hfq and repressing the synthesis of the abundant *lpp* mRNA (Guo *et al*, 2014). The results presented here show that MicL is cleaved by RNase E, and that precise cleavage is dependent on the presence of two adjacent hairpin structures 3′ to the cleavage site. Cleavage was found to be affected by both the stability of the two hairpins and the distance between them, with less impact of the sequence of the two hairpins or the sequence in and around the cleavage site. We also observed that the first stem-loop dictated the position of cleavage, and that Hfq promoted the specific cleavage. These results suggest that RNase E in conjunction with Hfq is recognizing helical structures leading to cleavage at a specified distance from this recognition element.

### sRNAs are differentially affected by *rne-3071*

The sequence of the MicL cleavage site is similar to the RNase E core motif identified in a recent study mapping transcriptome-wide RNase E cleavage sites via transient inactivation of RNase E followed by high-throughput RNA sequencing in *S. enterica* (Chao *et al*, 2017). However, even in this high-throughput study, the RNA levels of MicL and MicL-S were not severely affected by the transient inactivation of the RNase E, at least compared to other known RNase E sRNA substrates, analogous to what we observed (Fig 1).

The RNase E temperature-sensitive allele (*rne-3071*) in *E. coli* has a C to T transition at nucleotide position 742 resulting in the substitution of a leucine for a phenylalanine at amino position 68, near the nucleotide binding motif (Apirion, 1978; McDowall *et al*, 1993). This temperature-sensitive mutation was found to increase the chemical half-life of total pulse-labeled RNA (Kuwano *et al*, 1977) and affect the stability of single RNase E substrates (Babitzke & Kushner, 1991). Our results show different degrees of sRNA cleavage for the *rne-3071* strain at the non-permissible temperature; cleavage is almost completely abolished for ArcZ, significantly decreased for CpxQ and only slightly decreased for MicL (Fig 1A). Nevertheless, we think MicL is in fact a substrate for RNase E since the extent of MicL processing is decreased upon RNase E depletion in vivo (Fig 1B) and cleavage is observed at the same position with the purified NTD of RNase E in vitro (1-529) (Fig 2).

Our observations are consistent with early studies of the known RNase E substrates RNAI and T4 mRNA, for which it was found that not all RNase E decay intermediates are reduced in the *rne-3071* strain at the non-permissive temperature (Ehretsmann *et al*, 1992; McDowall *et al*, 1994). In fact, when comparing RNAI cleavage by wild-type and *rne-3071* at the non-permissive temperature, several decay products were similar between the two strains while some products were found to be enhanced and others were decreased. Furthermore, the *rne-3071* strain seemed to generate new RNAI decay bands not seen in the *rne*^+^ strain at the non-permissive temperature, leading the authors to warn about assigning RNase E cleavage sites based entirely on changes in RNA abundance in the *rne-3071* strain (McDowall *et al*, 1994).

### Recognition of MicL stem-loops by RNase E

From our mutational analyses, we propose the structural elements at the 3′ end of the MicL RNA are being recognized by RNase E in what is an “internal entry” pathway for cleavage. The 5′ end of in vivo transcribed MicL RNA is triphosphorylated (Thomason *et al*, 2015) and thus is less likely to recognized by RNase E. It is interesting to note that several known RNase E substrates have two stem-loop structures either preceding or following the cleavage site. This is the case for 9S RNA where two stem-loops are predicted to precede the single-stranded RNase E cleavage site and disruptions of these structures perturb cleavage while sequences 3′ of the cut site are dispensable (Cormack & Mackie, 1992). Similarly, three closely spaced stem-loop structures succeed the RNase E cleavage site of RNAI, while 5′ sequences are dispensable (McDowall et al, 1994). Interestingly, although the predominant cleavage site occurs 4 nucleotides from the first 3’ stem-loop structure of RNAI, a less intense cleavage site also occurs 4 nucleotides upstream of the second hairpin. Thus, it would seem, like with MicL, two closely spaced hairpins dictate cleavage of RNAI at a specified distance from an adjacent hairpin.

We suggest that the regions of RNase E observed to interact with helical structures of RprA and SdsR (Bandyra *et al*, 2018) in the crystal structures comprise the domain responsible for structural recognition of MicL. Consistent with this view the RNase E 1-529 8X mutant did not cleave Δ222 in vitro (Fig 8B). Due to the tetrameric organization of RNase E, it is conceivable that the two MicL 3′stem-loops can contact two adjacent subunits within the principle dimer, and that these close contacts anchor the single stranded cleavage site near the active site of the protein, explaining the requirement for a specific distance between the two 3′stem-loops as well as the specific site of cleavage relative to the first stem-loop. Given that MicL is a known Hfq binding sRNA (Guo *et al*, 2014) and Hfq likely interacts with the rho-independent terminator structure (Otaka *et al*, 2011; Sauer & Weichenrieder, 2011), an alternate explanation for the distance requirement is the need for a ternary complex between MicL, RNase E and Hfq. In this model, RNase E could bind the first stem-loop, while Hfq binds the second stem-loop. The lack of Δ222 cleavage by RNase E 1-529 when Hfq is omitted in vitro could be explained by this hypothesis.

### RNase E recognition of a combination of sRNA features could lead to a cleavage hierarchy

Other 3′ UTR-derived sRNAs that are cleaved by RNase E are predicted to have two closely spaced stem-loops at their 3′ ends, the second one corresponding to the Rho-independent transcription terminator (Chao et al, 2012; Chao & Vogel, 2016). Thus, we predict that structural features like what we have found for MicL help direct cleavage of other RNAs, especially 3′ UTR-derived sRNAs. We note that features in addition to the stem-loops could contribute to the very specific cleavage such that an sRNA might be recognized by a combination of 5′ sensing and ‘internal entry’. The individual recognition elements—5′ end, single stranded AU-rich sequence, 3′ stem loops and Hfq binding—each could contribute differently to the efficiency of 3′ UTR sRNA cleavage. This possibility is underscored by the differences in the sensitivities of MicL and CpxQ to the *rne-3017* allele and RNase E depletion. Overall our results suggest that RNase E substrate recognition is far more nuanced than initially imagined, allowing for even more levels or regulation and a hierarchy of RNase E-mediated cleavage.

## Materials and Methods

### Bacterial strains and plasmids

The bacterial strains and plasmids used in the study are listed in Appendix Tables S1 and S2, respectively. For plasmid construction, the desired gene fragments were either cloned into the vectors using the Gibson Assembly Cloning Kit (New England Biolabs) or were generated by PCR amplification using MG1655 genomic DNA as a template and, after digestion with restriction enzymes, were cloned into the corresponding sites of the indicated vectors. All oligonucleotides used are listed in Appendix Table S3. pBR* (pMSG14, (Guo *et al*, 2014)) is a derivative of the pBR322-derived pBRplac vector (here denoted as pBR, (Guillier & Gottesman, 2006)) in which the ampicillin cassette was replaced by the kanamycin cassette. All cloning was performed using *E. coli* TOP10 cells (Invitrogen), and all mutations and plasmid inserts were confirmed by sequencing. Mfold (Zuker, 2003) was used to predict the structures of the RNAs expressed from the constructs.

### Growth conditions

Unless indicated otherwise, strains were grown aerobically at 37°C in LB (10 g tryptone, 5 g yeast extract, 5 g NaCl per L) to an OD_600_ of 1.0. The following compounds were added at the following final concentrations where indicated; IPTG at 1 mM, ampicillin at 100 µg/ml, kanamycin at 30 µg/ml, chloramphenicol at 25 µg/ml and tetracycline at 12.5 µg/ml.

To deplete RNase E, *E. coli* strain KSL2000, with the pBAD-RNE plasmid (Lee *et al*, 2002), was grown overnight at 37°C in LB supplemented with ampicillin, chloramphenicol, tetracycline and 0.1 % arabinose. Strains without arabinose failed to grow. The overnight culture (100 μl) diluted into 30 ml of LB supplemented with ampicillin, chloramphenicol, tetracycline and 0.1% arabinose and grown at 37°C to and OD_600_ of 0.5. The culture was then washed 3X with 40 ml of LB media and resuspended in LB with ampicillin, chloramphenicol, tetracycline to an OD_600_ of 0.2. The culture was then split into two 15 ml-cultures with one culture having 0.1% arabinose and the other 0.1% glucose. The two cultures were then grown at 37°C to OD_600_ ~1 and ~1.2, and 1 ml of each culture was harvested and subject to total RNA extraction and northern analysis as described below. As expected, the culture with glucose grew slower than the culture with arabinose.

### Northern analysis

For northern analysis, total RNA was extracted with TRIzol Reagent as described (Hao *et al*, 2016). Briefly, 1 to 5 ml cells grown to OD600 ~1.0 (or indicated otherwise) were collected and resuspended in 1 ml of TRIzol^TM^ Reagent (Thermo Fisher Scientific) with repetitive pipetting to lyse the cells. The mixture was incubated at room temperature for 5 min, 0.2 ml of chloroform was added, and the sample was vortexed vigorously. The sample was then centrifuged for 15 min at 4°C at 11,000 rpm. The top ~0.6 ml of the aqueous phase was transferred to a new Eppendorf tube and 0.5 ml of isopropyl alcohol added. After 10 min incubation at room temperature the sample was centrifuged at 11,000 rpm for 5 min at 4°C. The precipitated RNA pellet was washed with 1 ml of 75% ethanol and finally air-dried and dissolved in DEPC treated dH2O. Total RNA concentration was determined based on OD_260_.

Northern blots were performed as described (Hao *et al*, 2016). Briefly, 10 μg of total RNA was separated on an 8% polyacrylamide–7M urea gel (USB Corporation) in 1X TBE and transferred to Zeta-Probe membrane (Bio-Rad) overnight at 20 V in 0.5X TBE. Oligonucleotides were end-labeled with γ-^32^P-ATP by T4 polynucleotide kinase (New England Biolabs). Membranes were UV cross-linked and hybridized overnight with the labeled probe at 45°C in UltraHyb (Ambion) hybridization buffer. Following hybridization, membranes were washed once with 2X SSC + 0.1% SDS followed by a 10-min incubation at 45°C with 2X SSC + 0.1% SDS. Membranes were subsequently washed 5X with 0.2X SSC + 0.1% SDS, allowed to air dry for 5 min, and exposed to KODAK Biomax X-ray film at −80°C.

### In vitro RNA synthesis

The MicL (308 nt), Δ222 (86 nt), Δ222-stemAΔ2 (72 nt), Δ222+10 (96 nt), Δ222+15 (101 nt) and 9S (243 nt) RNAs were synthesized using a MEGAshortscript T7 Transcription kit (Ambion) using manufacturer’s guidelines. Purified RNA was quantified by OD_260_ measurements. For experiments that called for 9S to be dephosphorylated, 20 pmol T7-transcribed RNA was treated with Calf Intestinal Phosphatase (CIP) (New England Biolabs) using manufactures guidelines and then extracted with phenol:chloroform:IAA and ethanol precipitated.

### In vitro assay of RNase E activity

The in vitro cleavage assays with the purified RNase E derivatives and in vitro transcribed RNAs were carried out with slight modifications of a previously reported protocol (Chao & Vogel, 2016). Briefly, the RNA was diluted in Buffer A (25 mM Tris pH 7.5, 50 mM NaCl, 50 mM KCl, 10 mM MgCl_2_, 1 mM DTT and 0.5 U µl^-1^ RNase Out (Thermo Fisher Scientific)), and heat denatured at 95°C for 1 min and cooled on ice for 5 min. In a 10 µL reaction, 300 nM RNA was incubated wiht 300 nM Hfq_6_ for 10 min at 30°C. Purified RNase E (1-529) or D26N,D28N,D338N mutant RNase E (1-529) or 8X mutant RNase E (1-529) (300 nM) was added and reactions incubated for an additional 30 min at 30°C. Ammonium acetate stop solution (Ambion) (7.5 µl) was then added along with 82.5 µl dH_2_O and 100 µL of phenol:chloroform:IAA. The aqueous phase was separated from the organic phase by PLG-tubes (5PRIME) and the RNA precipitated from the aqueous phase by the addition of 10 µL of 3M sodium acetate, 250 µl of 100% ethanol and 3 µL of glycogen and incubation at −80°C for 30 min. The RNA was then centrifuged at 15,000 rpm for 30 min at 4°C, and the pellet washed with 1 ml of 70 % ethanol and resuspended in 20 µL of Gel Loading Buffer II (Ambion). An aliquot (10 µL) was analyzed via northern analysis as described above.

## Acknowledgements

We thank B. Luisi for helpful discussions and the kind gifts of the purified 1-529 domains of wild type, D26N,D28N,D338N mutant and 8X mutant RNase E. We are grateful to K. Lee for sharing the RNase E depletion strain. We also thank S. Gottesman and B. Luisi for comments on this manuscript. Work carried out by TBU, ABK and GS was supported by the Intramural Research program of the *Eunice Kennedy Shriver* National Institute of Child Health and Human Development. Work carried out by KJB was supported by the Wellcome Trust.

## References

Apirion D (1978) Isolation, genetic mapping and some characterization of a mutation in *Escherichia coli* that affects the processing of ribonuleic acid. Genetics 90: 659–671

Babitzke P, Kushner SR (1991) The Ams (altered mRNA stability) protein and ribonuclease E are encoded by the same structural gene of Escherichia coli. Proc Natl Acad Sci USA 88: 1–5

Bandyra KJ, Bouvier M, Carpousis AJ, Luisi BF (2013) The social fabric of the RNA degradosome. Biochim Biophys Acta 1829: 514–522

Bandyra KJ, Wandzik J, Luisi BF (2018) Substrate recognition and auto-inihibition in the central ribonuclease RNase E. Mol Cell **Under revision.**

Chao Y, Li L, Girodat D, Förstner KU, Said N, Corcoran C, Śmiga M, Papenfort K, Reinhardt R, Wieden HJ, Luisi BF, Vogel J (2017) In vivo cleavage map illuminates the central role of RNase E in coding and non-coding RNA pathways. Mol Cell 65: 39–51

Chao Y, Papenfort K, Reinhardt R, Sharma CM, Vogel J (2012) An atlas of Hfq-bound transcripts reveals 3’ UTRs as a genomic reservoir of regulatory small RNAs. EMBO J 31: 4005–4019

Chao Y, Vogel J (2016) A 3’ UTR-derived small RNA provides the regulatory noncoding arm of the inner membrane stress response. Mol Cell 61: 352–363

Cherepanov PP, Wackernagel W (1995) Gene disruption in *Escherichia coli*: TcR and KmR cassettes with the option of Flp-catalyzed excision of the antibiotic-resistance determinant. Gene 158: 9–14

Clarke JE, Kime L, Romero AD, McDowall KJ (2014) Direct entry by RNase E is a major pathway for the degradation and processing of RNA in *Escherichia coli*. Nucleic Acids Res 42: 11733–11751

Cormack RS, Mackie GA (1992) Structural requirements for the processing of *Escherichia coli* 5 S ribosomal RNA by RNase E in vitro. J Mol Biol 228: 1078–1090

Court DL, Swaminathan S, Yu D, Wilson H, Baker T, Bubunenko M, Sawitzke J, Sharan SK (2003) Mini-l: a tractable system for chromosome and BAC engineering. Gene 315: 63–69

Datsenko KA, Wanner BL (2000) One-step inactivation of chromosomal genes in *Escherichia coli* K-12 using PCR products. Proc Natl Acad Sci USA 97: 6640–6645

Deana A, Celesnik H, Belasco JG (2008) The bacterial enzyme RppH triggers messenger RNA degradation by 5’ pyrophosphate removal. Nature 451: 355–358

Del Campo C, Bartholomäus A, Fedyunin I, Ignatova Z (2015) Secondary structure across the bacterial transcriptome reveals versatile roles in mRNA regulation and function. PLoS Genet 11: e1005613

Ehretsmann CP, Carpousis AJ, Krisch HM (1992) Specificity of *Escherichia coli* endoribonuclease RNase E: in vivo and in vitro analysis of mutants in a bacteriophage T4 mRNA processing site. Genes Dev 6: 149–159

Göpel Y, Khan MA, Görke B (2016) Domain swapping between homologous bacterial small RNAs dissects processing and Hfq binding determinants and uncovers an aptamer for conditional RNase E cleavage. Nucleic Acids Res 44: 824–837

Guillier M, Gottesman S (2006) Remodelling of the Escherichia coli outer membrane by two small regulatory RNAs. Mol Microbiol 59: 231–247

Guo MS, Updegrove TB, Gogol EB, Shabalina SA, Gross CA, Storz G (2014) MicL, a new σE-dependent sRNA, combats envelope stress by repressing synthesis of Lpp, the major outer membrane lipoprotein. Genes Dev 28: 1620–1634

Hao Y, Updegrove TB, Livingston NN, Storz G (2016) Protection against deleterious nitrogen compounds: role of σ^S^-dependent small RNAs encoded adjacent to *sdiA*. Nucleic Acids Res 44: 6935–6948

Kaberdin VR (2003) Probing the substrate specificity of Escherichia coli RNase E using a novel oligonucleotide-based assay. Nucleic Acids Res 31: 4710–4716

Kim HM, Shin JH, Cho YB, Roe JH (2014) Inverse regulation of Fe- and Ni-containing SOD genes by a Fur family regulator Nur through small RNA processed from 3’UTR of the *sodF mRNA*. Nucleic Acids Res 42: 2003–2014

Kuwano M, Ono M, Endo H, Hori K, Nakamura K, Hirota Y, Ohnishi Y (1977) Gene affecting longevity of messenger RNA: a mutant of *Escherichia coli* with altered mRNA stability. Mol Gen Genet 154: 279–285

Lee K, Bernstein JA, Cohen SN (2002) RNase G complementation of rne null mutation identifies functional interrelationships with RNase E in *Escherichia coli*. Mol Microbiol 43: 1445–1456

Mackie GA (2013) RNase E: at the interface of bacterial RNA processing and decay. Nat Rev Microbiol 11: 45–57

McDowall KJ, Hernandez RG, Lin-Chao S, Cohen SN (1993) The ams-1 and rne-3071 temperature-sensitive mutations in the *ams* gene are in close proximity to each other and cause substitutions within a domain that resembles a product of the *Escherichia coli mre* locus. J Bacteriol 175: 4245–4249

McDowall KJ, Lin-Chao S, Cohen SN (1994) A+U content rather than a particular nucleotide order determines the specificity of RNase E cleavage. J Biol Chem 269: 10790–10796

Miyakoshi M, Chao Y, Vogel J (2015) Cross talk between ABC transporter mRNAs via a target mRNA-derived sponge of the GcvB small RNA. EMBO J 34: 1478–1492

Mohanty BK, Kushner SR (2016) Regulation of mRNA decay in bacteria. Annu Rev Microbiol 70: 25–44

Ono M, Kuwano M (1979) A conditional lethal mutation in an *Escherichia coli* strain with a longer chemical lifetime of messenger RNA. J Mol Biol 129: 343–357

Otaka H, Ishikawa H, Morita T, Aiba H (2011) PolyU tail of rho-independent terminator of bacterial small RNAs is essential for Hfq action. Proc Natl Acad Sci USA 108: 13059–13064

Sauer E, Weichenrieder O (2011) Structural basis for RNA 3’-end recognition by Hfq. Proc Natl Acad Sci USA 108: 13065–13070

Thomason MK, Bischler T, Eisenbart SK, Förstner KU, Zhang A, Herbig A, Nieselt K, Sharma CM, Storz G (2015) Global transcriptional start site mapping using differential RNA sequencing reveals novel antisense RNAs in *Escherichia coli*. J Bacteriol 197: 18–28

Yu D, Ellis HM, Lee EC, Jenkins NA, Copeland NG, Court DL (2000) An efficient recombination system for chromosome engineering in *Escherichia coli*. Proc Natl Acad Sci USA 97: 5978–5983

Zuker M (2003) Mfold web server for nucleic acid folding and hybridization prediction. Nucleic Acids Res 31: 3406–3415

